## SupplementaryMaterials for "Self-association enhances early attentional selection through automatic prioritization of socially salient signals"

**Supporting information**

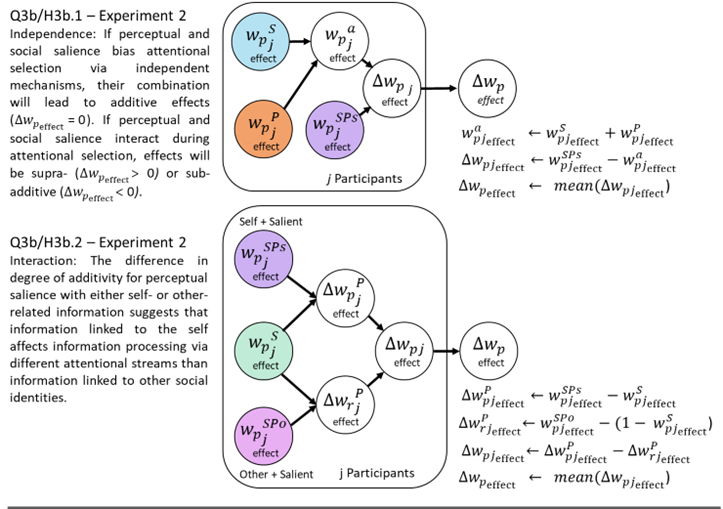

**Supplemental figure S1:** Formalization of interaction effect assessment, using attentional weights. Note that, instead of attentional weights, we report processing rates. However, the same effects that are reported in the main text are reproduced when attentional weights are used in the analysis.

**S2. Randomization and counterbalancing of within-task parameters**

Within the TOJ task, the following parameters were randomized (r) or counterbalanced (c) within and across blocks:

- Target hemifield: Whether the probe is shown left or right. (c)
- Stimulus location: Which out of 4 possible locations in each hemifield the probe is presented. Probe and reference are symmetrically aligned to keep the distance to fixation equal for each trial. (c)
- Response label position: Whether the probe label is presented above or below fixation. Probe and reference labels are always presented opposite each other vertically. (c)
- Order of SOAs: Whether, and by which amount, the probe precedes or follows the presentation of the reference. (r)
- Choice of probe: Which stimulus is defined as the probe in perceptual salience & perceptual baseline condition on each trial (r). The probe is determined by the shape for the social salience and social baseline conditions.

Within the matching task, the following parameters were randomized (r) or counterbalanced (c) within and across blocks:

- Condition order: Which stimulus pair (identity, matching) is shown next (r)
- Stimulus location: Whether the shape is shown above or below fixation. The label is always shown in the vertically opposite position (c)
- Stimulus frequency: Whether each identity and matching condition is shown with the same frequency within and across blocks (c)

**S3. Individual response data**

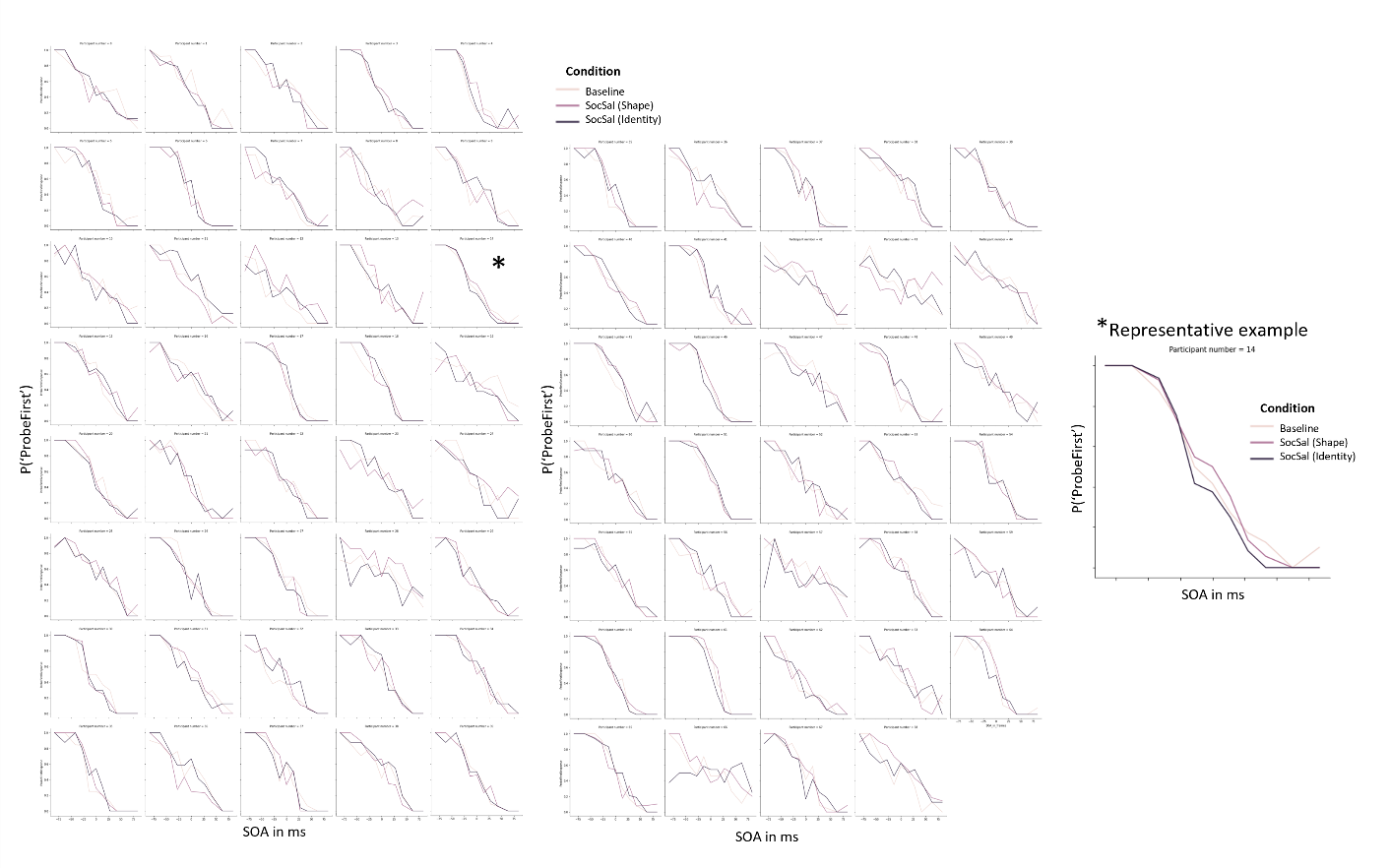

**Supplemental Figure S2:** Individual participant response data indicating the proportion with which participants responded that the probe flickered first as a function of stimulus onset (flicker) asynchrony. Different conditions are shown in different shadings. Panel on the right shows an example participant with a group-representative response pattern: an increase of ‘probe first’ responses when there the probe was self-associated and the shape of the stimulus had to be reported. This pattern is specifically visible at low onset asynchronies. Furthermore, this participant shows a decreased proportion of ‘probe first’ responses when the probe was self-associated and the social identity had to be reported.

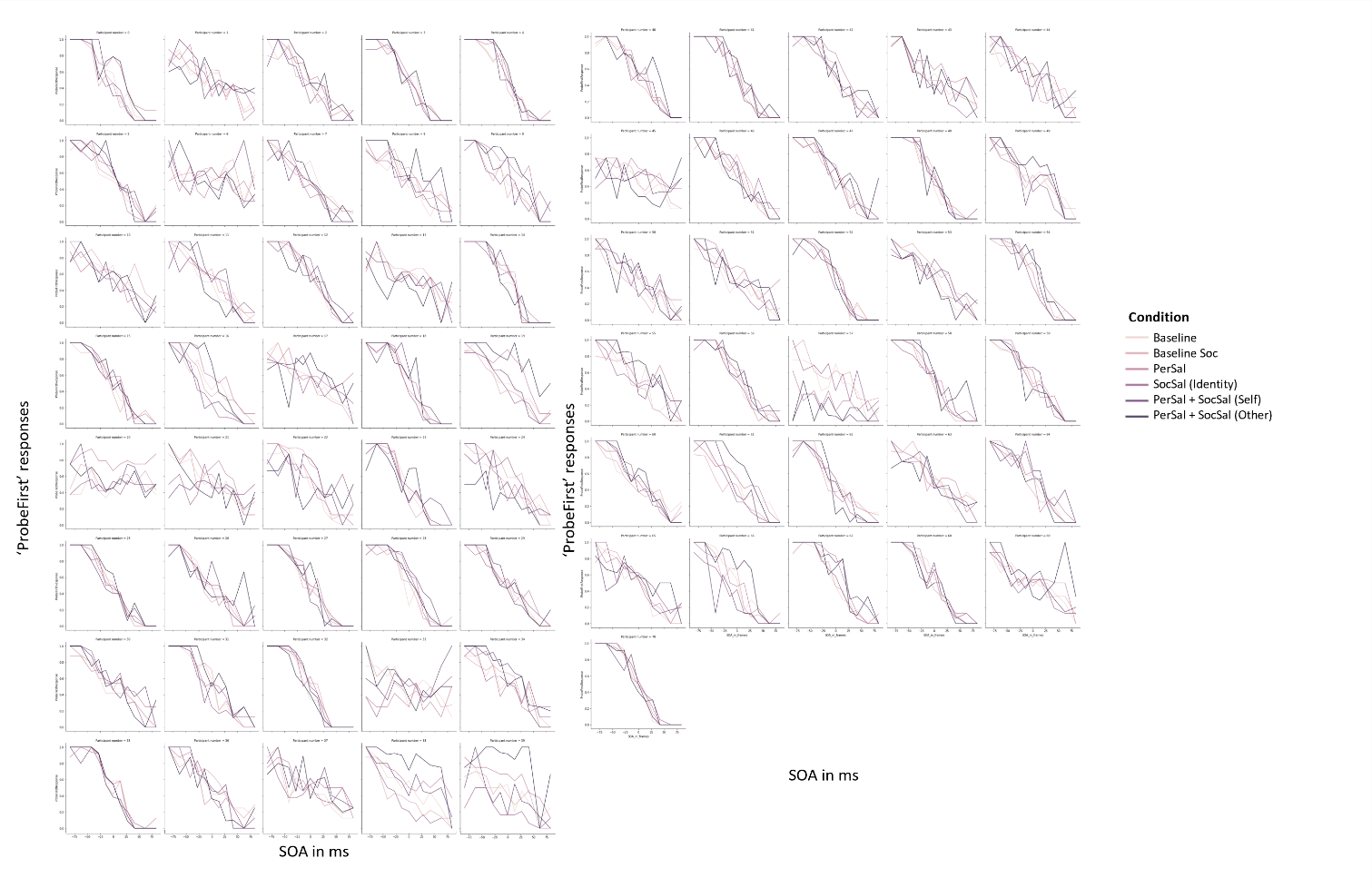

**Supplemental Figure S3:** Individual participant response data indicating the proportion with which participants responded that the probe flickered first as a function of stimulus onset (flicker) asynchrony. Different conditions are shown in different shadings.

**S4. Individual parameter estimates**

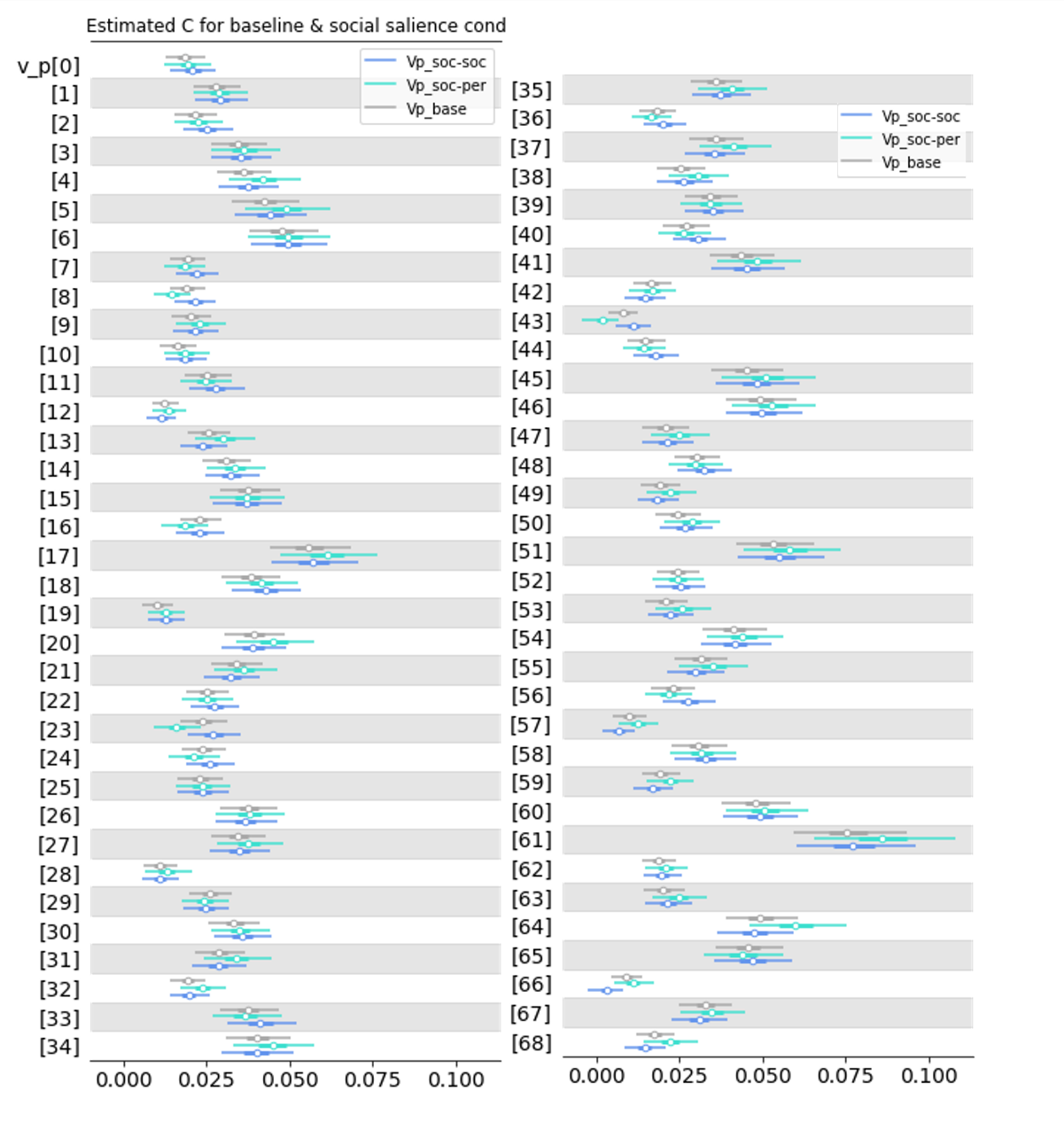
**Supplemental Figure S4:** Absolute processing rates (*v_p_*) for the probe stimulus (self-associated), shown for individual participants in Experiment 1. Different colours indicate different conditions: the baseline (*grey*; *v_p__base*), the social salience condition in which the shape had to be reported (*light blue*; *v_p__soc-per*) and the social salience condition in which the identity had to be reported (*dark blue*; *v_p__soc-soc*). Points show the estimated posterior means, thick lines indicate the central quartiles, and thin lines indicate the 95% highest density interval. Processing rates are given in items/ms.

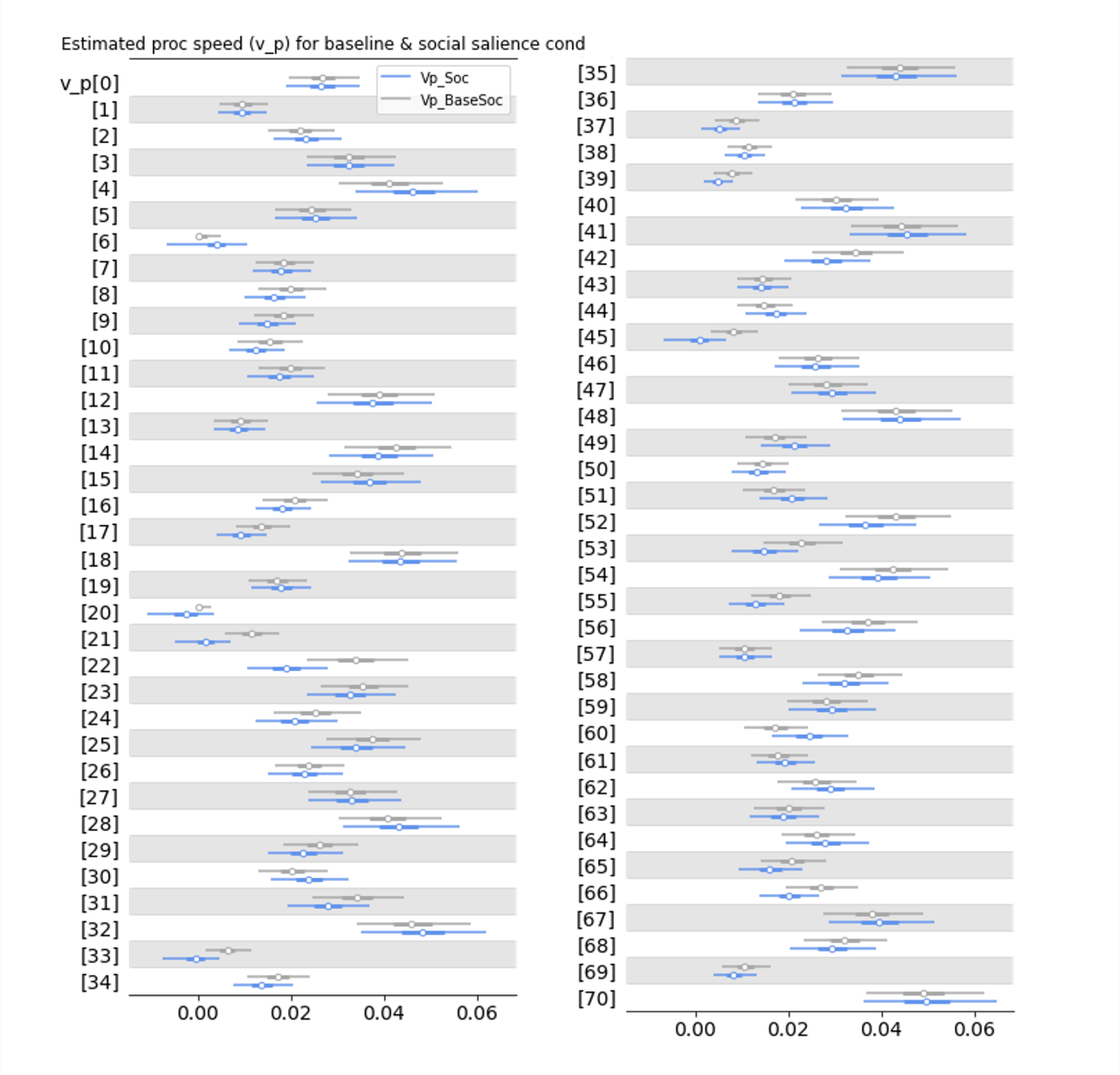

**Supplemental Figure S5:** Absolute processing rates (*v_p_*) for the probe stimulus (self-associated), shown for individual participants for the social baseline (*grey*; *v_p__baseSoc*) and the social salience condition in which the identity had to be reported (*dark blue*; *v_p__soc*). Points show the estimated posterior means, thick lines indicate the central quartiles, and thin lines indicate the 95% highest density interval. Processing rates are given in items/ms.

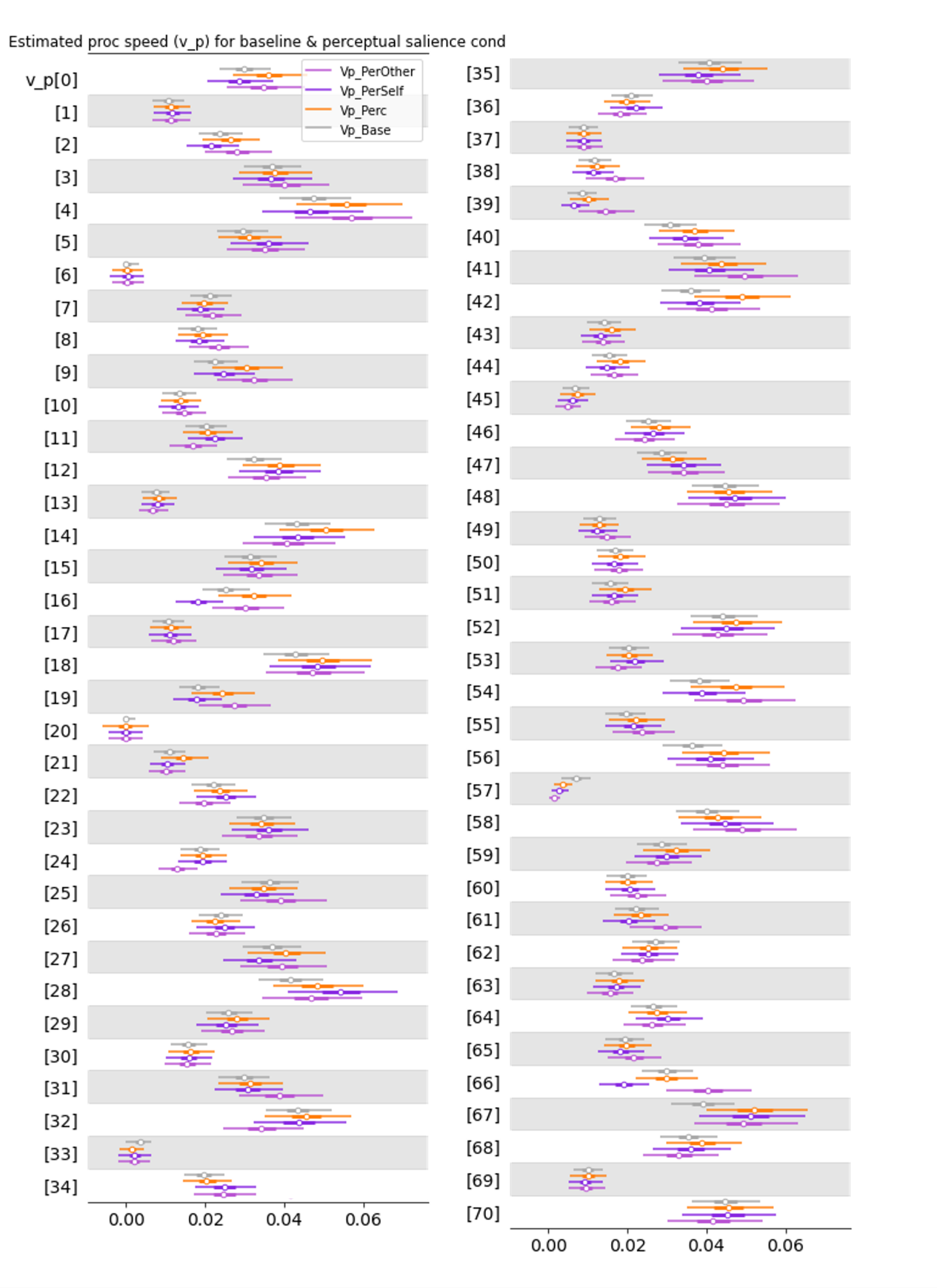

**Supplemental Figure S6:** Absolute processing rates (*v_p_*) for the probe stimulus (perceptually salient), shown for individual participants for the baseline (*grey*; *v_p__base*), for the mere perceptual salience condition (*orange*; *v_p__perc*), for the perceptual salience condition in which the probe was self-associated (*purple*; *v_p__perSelf*), and the perceptual salience condition in which the probe was other-associated (*pink*; *v_p__perOther*). Points show the estimated posterior means, thick lines indicate the central quartiles, and thin lines indicate the 95% highest density interval. Processing rates are given in items/ms.
